# Identification of putative effector genes across the GWAS Catalog using molecular quantitative trait loci from 68 tissues and cell types

**DOI:** 10.1101/808444

**Authors:** Cong Guo, Karsten B. Sieber, Jorge Esparza-Gordillo, Mark R. Hurle, Kijoung Song, Astrid J. Yeo, Laura M. Yerges-Armstrong, Toby Johnson, Matthew R. Nelson

## Abstract

Identifying the effector genes from genome-wide association studies (GWAS) is a crucial step towards understanding the biological mechanisms underlying complex traits and diseases. Colocalization of expression and protein quantitative trait loci (eQTL and pQTL, hereafter collectively called “xQTL”) can be effective for mapping associations to genes in many loci. However, existing colocalization methods require full single-variant summary statistics which are often not readily available for many published GWAS or xQTL studies. Here, we present PICCOLO, a method that uses minimum SNP p-values within a locus to determine if pairs of genetic associations are colocalized. This method greatly expands the number of GWAS and xQTL datasets that can be tested for colocalization. We applied PICCOLO to 10,759 genome-wide significant associations across the NHGRI-EBI GWAS Catalog with xQTLs from 28 studies. We identified at least one colocalized gene-xQTL in at least one tissue for 30% of associations, and we pursued multiple lines of evidence to demonstrate that these mappings are biologically meaningful. PICCOLO genes are significantly enriched for biologically relevant tissues, and 4.3-fold enriched for targets of approved drugs.

---

GWAS have discovered thousands of genetic associations with complex traits and diseases^1^. Most associations cannot be explained by protein coding changes and are expected to be driven by gene regulatory mechanisms^1,2^. Recent studies have shown that xQTLs explain a substantial proportion of complex trait heritability^3,4^. However, it is not enough to demonstrate that a GWAS index SNP is a statistically significant xQTL to conclude that the GWAS and xQTL associations are likely explained by the same underlying functional variant^5^. This requires that the two associations colocalize. A popular colocalization method developed by Giambartolomei *et. al*. determines if two associations are driven by the same signal using full single-variant summary statistics for the genomic region of interest^5^. Already, colocalization analyses have identified candidate effector genes for a variety of complex diseases and traits^6-9^. However, broader application of the colocalization approach is limited by the lack of readily available complete summary statistics from many published GWAS and xQTL studies. Even when the data are available, obtaining and harmonizing results takes significant effort. To overcome the need for full summary statistics and to expand the pool of available GWAS and xQTL studies for colocalization, we developed PICCOLO, a colocalization test using Probabilistic Identification of Causal SNPs (PICS) credible sets that can be estimated using only an index SNP p-value and a linkage disequilibrium reference panel^10^. Using PICCOLO, we colocalized all associations in the NHGRI-EBI GWAS Catalog^11^ with eQTLs from The Genotype-Tissue Expression (GTEx) Project^12^ along with xQTLs from 27 additional studies^13-38^ (Supplementary Table 1).

We created the PICCOLO algorithm by adapting the coloc method^5^ to use PICS for estimating causal SNP probabilities. Once causal SNP probabilities for both genetic associations have been estimated, PICCOLO performs a statistical test to evaluate their overlap. In contrast to coloc (Fig. 1a), PICCOLO enables the assessment of colocalization for any two genetic signals using only published index SNPs (Fig. 1b). PICS estimates the posterior probability of causality for each SNP within a locus using the LD structure from a reference data set and strength of association^10^. Therefore, all that is required to generate posterior probabilities with PICS is the index SNP identifier, the p-value of the index SNP, and the ancestry of the study population. As with coloc, PICCOLO calculates the posterior probability that two genetic associations are either shared (H4) or not shared (H3). In contrast to coloc, PICCOLO does not test the hypotheses of no association for either trait (H1, H2), or both traits (H0).

**Figure 1:**
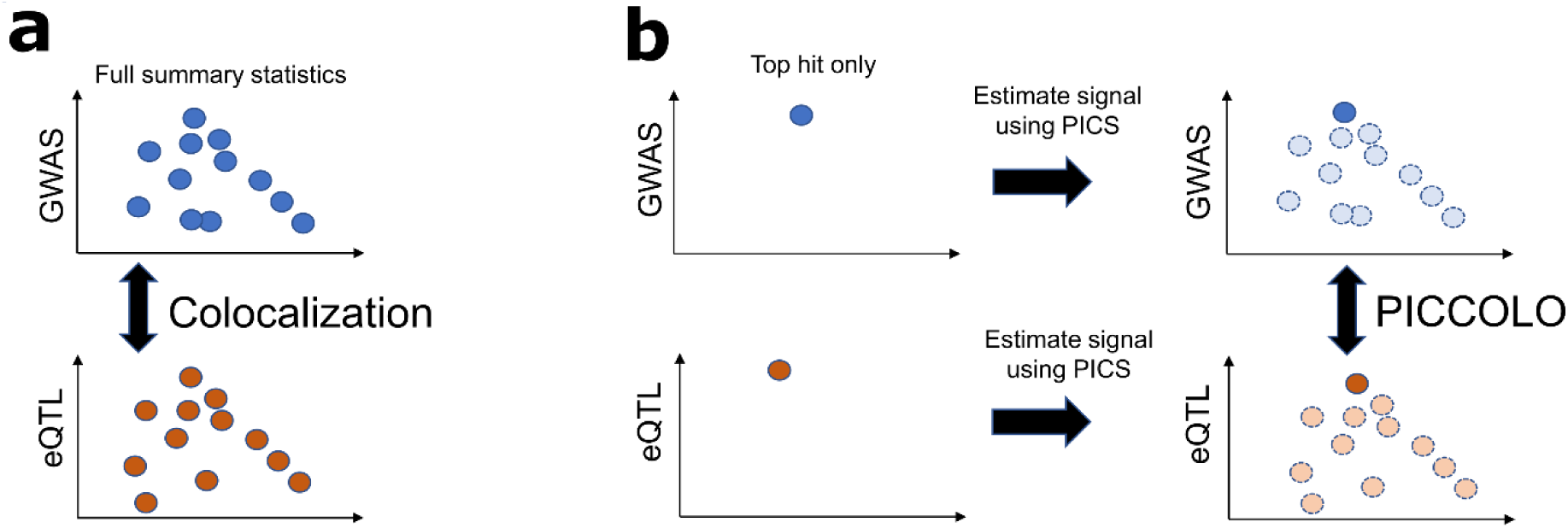
Colocalization of genetic signals using PICCOLO. **(a)** Colocalization requires full summary statistics from both the GWAS and xQTL studies. **(b)** PICCOLO uses association top hits and estimates the “missing” data using PICS. The PICS sets are then colocalized using the *coloc* method.

To evaluate the performance of PICCOLO, we compared PICCOLO with coloc results for 1,490 genome-wide significant loci (P ≤ 5 × 10^−8^) across 13 diverse traits analyzed in UK Biobank^39^ (Supplementary Table 2). These loci were tested using coloc and PICCOLO across the 44 GTEx Version 6p eQTL tissues (Supplementary Data 1). Coloc analyses were conducted on full variant-level summary statistics across each locus for both the GWAS and GTEx results. PICCOLO analyses were conducted with just the most significant GWAS and GTEx SNP in each locus. In comparing the two methods, true positives were defined as coloc genes with a posterior probability of H4 ≥ 0.8. Analysis of the PICCOLO parameters identified that PICCOLO H4 ≥ 0.9 and an xQTL P ≤ 1 × 10^−5^ are parameters that provide a strong predictive power for coloc (positive predictive value = 0.89) and reasonable sensitivity (0.52) of coloc results (Fig. 2, Supplementary Figs. 1-3, see methods). We did not observe any impact of allele frequency or credible set size on PICCOLO performance (Supplementary Figs. 4-6). Therefore, differences between PICCOLO and coloc are likely due to limitations of the PICS estimations from the index SNPs.

**Figure 2:**
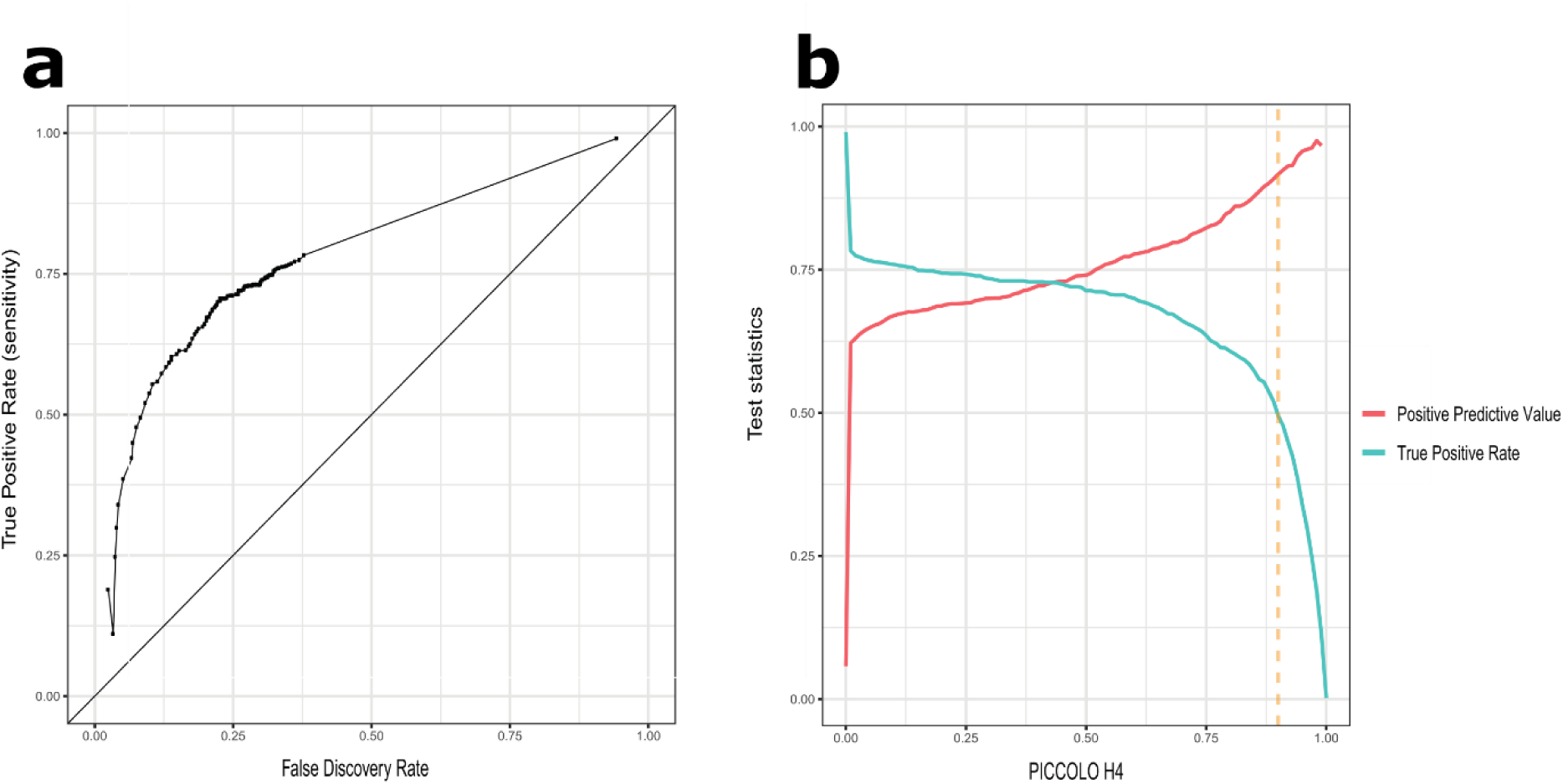
Calibration of PICCOLO with coloc. Using index SNPs, PICCOLO was evaluated for the ability to predict colocalization (coloc H4 ≥ 0.80). **(a)** ROC comparing PICCOLO to coloc in a tissue agnostic manner. **(b)** Using the balanced PICCOLO parameters (orange dashed line, H4 ≥ 0.90 and an xQTL P ≤ 1 × 10^−5^), PICCOLO has a positive predictive value (red line) of 0.89 and sensitivity (blue line) of 0.52 for coloc (coloc H4 ≥ 0.8).

Since PICCOLO estimates colocalization of genetic associations without complete summary statistics, we used PICCOLO to identify possible causal genes using results available in the NHGRI-EBI GWAS Catalog^11^. We generated PICS probabilities for 23,012 genome-wide significant associations and 23,353 non-genome-wide significant associations. In addition, we generated xQTL PICS probabilities for 44 tissues in GTEx V6p (166,987 unique index SNPs) and 27 other studies (148,259 unique index SNPs)^13-38^. In total, we used xQTLs from 32 broad tissue groups, representing 68 tissues and cell types (Supplementary Table 3).

PICCOLO was then used to assess colocalization of the GWAS and xQTL associations (Supplementary Data 2). Applying our previously selected parameters (H4 ≥ 0.9, xQTL P ≤ 1 × 10^−5^), we found 6,628 (29%) genome-wide significant associations to have ≥ 1 PICCOLO colocalization (Supplementary Data 2). Of the 6,628 associations with PICCOLO colocalizations, GTEx eQTLs uniquely accounted for colocalizations for 2,500 associations (39%), non-GTEx xQTLs accounted for 1,802 associations (27%), and 2,240 (33%) associations were accounted for by both. These results highlight the added value of using index SNPs across additional xQTL sources to map genetic associations to putative effector genes.

Of GWAS loci with PICCOLO colocalizations, 2,730 (62%) colocalized with one gene in one or more cell types (gene-specific loci). Of those, 1,189 (27%) had colocalization with a single gene in a single cell type (gene- and tissue-specific loci, Fig. 3a). These results suggest that the majority of GWAS loci are driven by a single effector gene (Fig. 3b). Next, we wanted to determine how tissue-specific colocalizations were. We grouped xQTL tissues into 32 broad tissue groups (Supplementary Table 3) and found that 44% of associations colocalized with xQTLs from single tissue groups (tissue-specific loci) (Fig. 3c). These findings indicate that PICCOLO can resolve 2,730 (12%) of all genome-wide significant associations to a single gene, and 1,839 (8%) of associations to a single tissue group.

**Figure 3:**
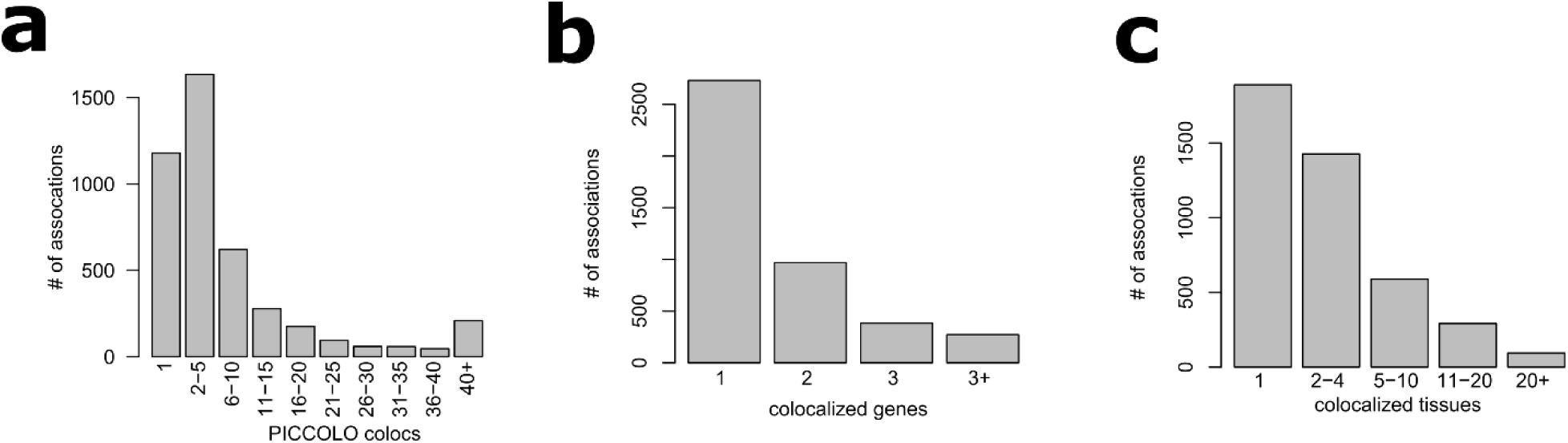
Colocalization of the NHGRI-EBI GWAS Catalog with xQTLs using PICCOLO. **(a)** Distribution of the number of colocalizations (tissue-gene pairs) at each GWAS association. **(b)** Distribution of the number of PICCOLO genes at each association. **(c)** Distribution of the number of tissue groups with colocalizations at each GWAS association.

Next, we hypothesized that PICCOLO genes that were specific to trait categories would be enriched for colocalizations in tissues that are biologically relevant. Using the Medical Subject Heading (MeSH) terms we grouped GWAS into 18 trait categories (Supplementary Data 3). For each trait category, we determined the number of PICCOLO genes that were specific or shared across each respective category (Fig. 4a). Blood traits had 1.5× more category specific genes than nonspecific genes (P = 4.7 × 10^−6^). In contrast, inflammatory traits had 2.2× more nonspecific genes than category specific genes (P = 0.02). While 2,383 (71%) of PICCOLO genes could be attributed to a single trait category, 66 (2%) were highly pleotropic, spanning > 4 trait categories (Fig. 4b, Supplementary Data 4). For each trait category, we tested for the enrichment of xQTL tissues for PICCOLO genes specific to that trait compared to all other PICCOLO colocalizations. Most enriched tissues were clearly biologically relevant. For example, cardiovascular traits were enriched for PICCOLO colocalizations in heart and blood vessel. In contrast, the uterus, vagina, prostate, and pituitary tissue enrichments observed in inflammation traits were less obvious, but biologically insightful (Fig. 4c). The enrichment of colocalizations in sex-specific tissues is consistent with established studies highlighting the strong sex bias and key role of sex hormones in autoimmunity^40-42^. Together, these results demonstrate that PICCOLO identifies genes that are likely specific to one trait category and in biologically relevant tissues.

**Figure 4:**
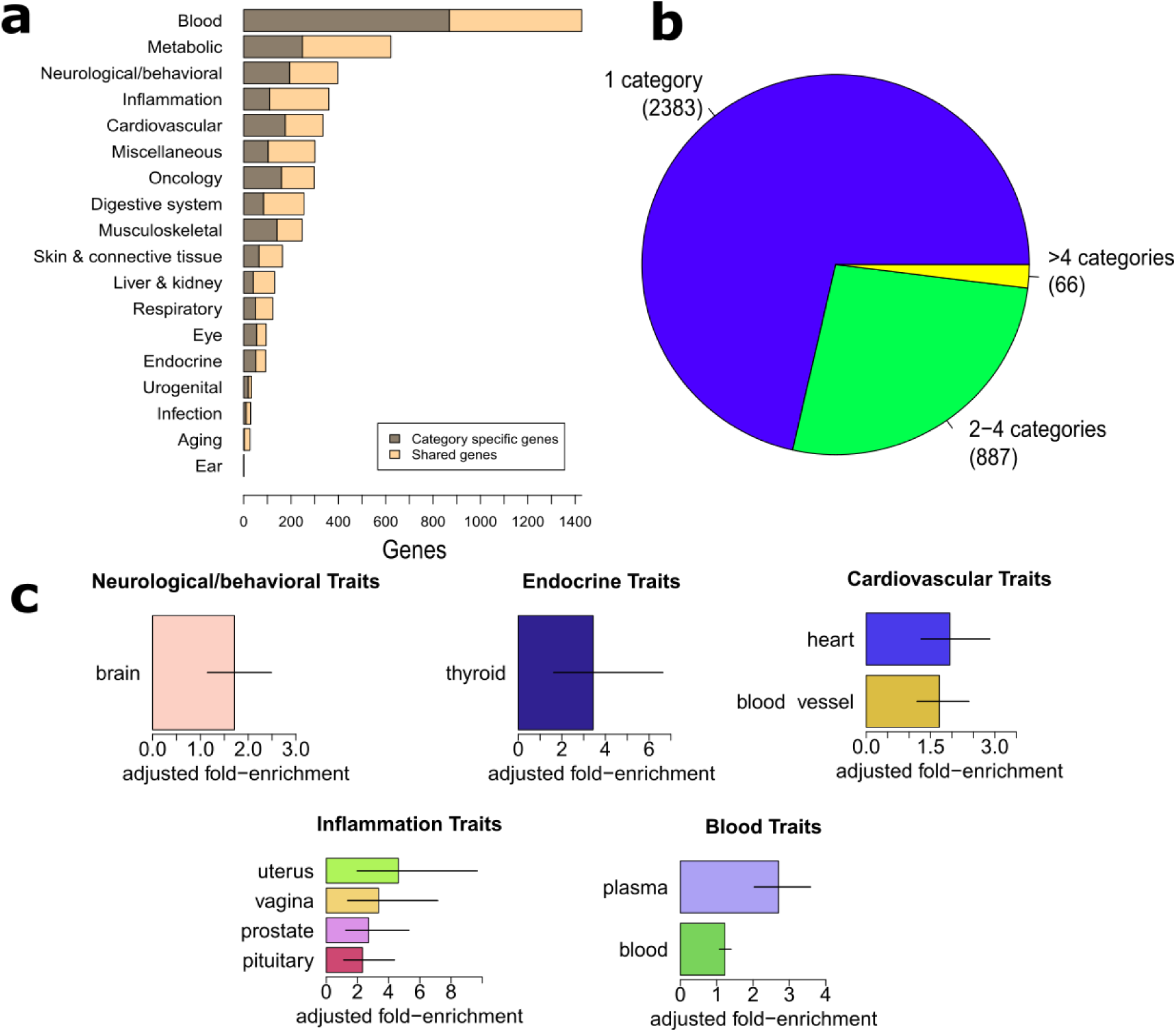
PICCOLO results across different trait categories. **(a)** Number of PICCOLO genes that are shared across one or more trait categories (beige) and number of PICCOLO genes that are specific to a trait category (grey). **(b)** Proportion of PICCOLO genes that map to a single therapy category (blue), 2-4 categories (green), and >4 categories (yellow). **(c)** Enrichment of QTL tissues for trait category specific PICCOLO genes across neurological/behavioral, endocrine, cardiovascular, inflammation, and blood-related traits. Bars show 95% confidence intervals. All enrichments are significant at an FDR of 0.5. If a trait category or tissue is not shown, it means that the category specific genes were not enriched for PICCOLO genes that colocalize in those tissues.

Next, we tested whether PICCOLO genes were more likely to be disease modulating. For our first analysis, we assessed the enrichment of PICCOLO genes among all genes implicated in rare diseases documented in the Online Mendelian Inheritance in Man (OMIM) Catalog^43^ (Supplementary Data 5). Both genes nearest to GWAS associations and non-colocalized genes showed significant 1.2-fold enrichments for OMIM genes (P = 2.4 × 10^−9^); however, PICCOLO genes were enriched 2.2-fold (P =1.4 × 10^−12^, Fig. 5 top). We observed similar enrichments when the analysis was limited to PICCOLO colocalizations with QTLs of a single gene and/or within a single tissue (OR = 2.3, P = 5.4 × 10^−17^). These data demonstrate that genes identified as influencing complex traits by colocalization are more likely to cause rare disease.

**Figure 5:**
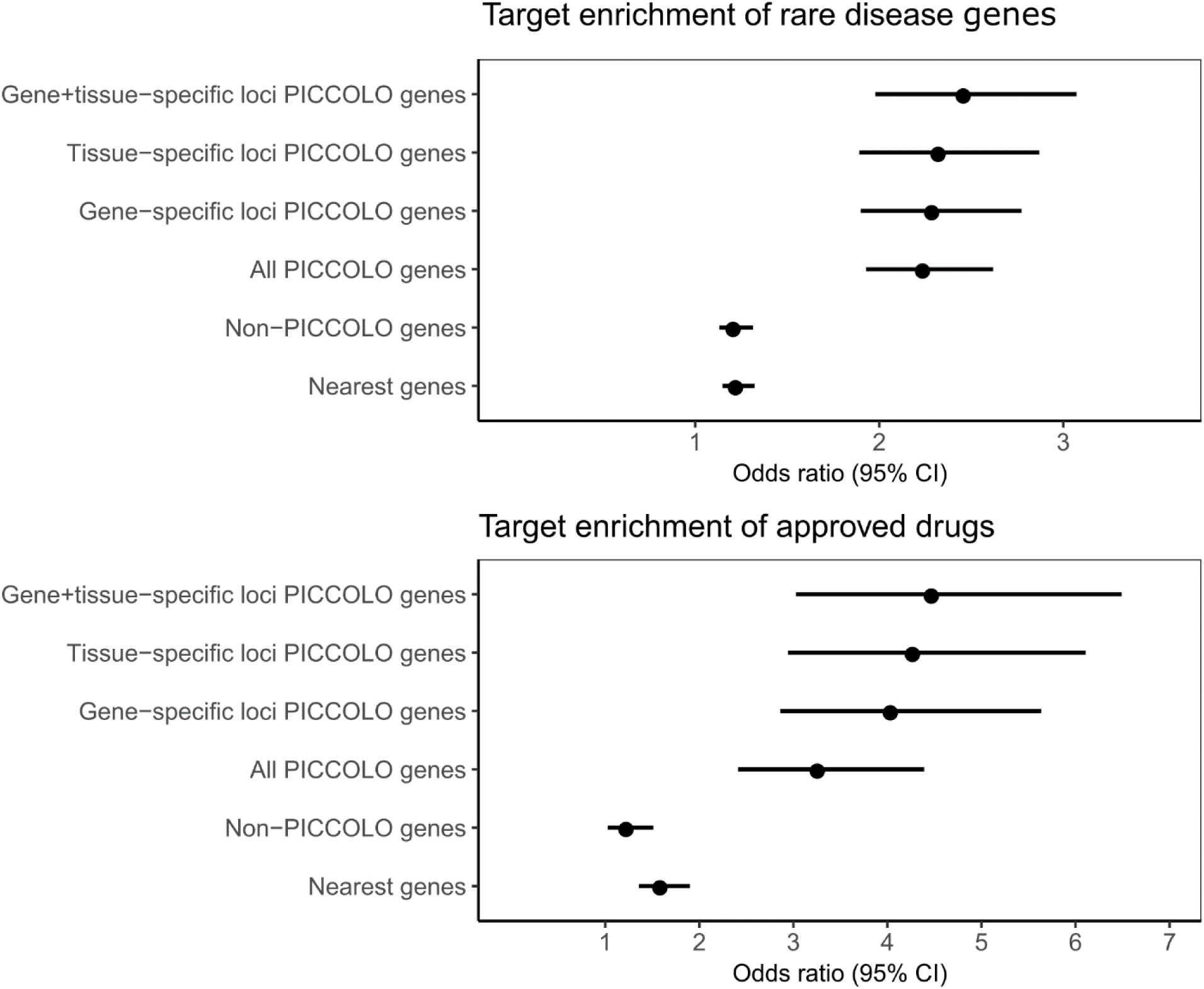
Enrichment of rare disease genes, approved drug targets and overlap between drug targets and PICCOLO genes. Enrichments for rare disease genes in OMIM (top), and approved drug targets in the United States and the European Union (bottom). Enrichment was tested for genes nearest to GWAS index SNPs (Nearest genes), genes that were found to not be colocalized using PICCOLO (Non-PICCOLO genes), all genes found to be colocalized using PICCOLO (All PICCOLO genes), PICCOLO genes within loci where only one gene is colocalized (Gene-specific loci PICCOLO genes), genes within loci where colocalization occur in a single tissue (Tissue-specific loci PICCOLO genes), and PICCOLO genes within loci where a single gene is colocalized in a single tissue (Gene+tissue-specific loci PICCOLO genes). Bars show 95% confidence intervals.

Lastly, we assessed enrichment of PICCOLO genes amongst the target genes of approved drugs in the Pharmaprojects database. Overall, these genes were 2.7-fold enriched (P = 1.4 × 10^−11^) for successful drug targets, compared to 1.2- and 1.6-fold enrichments for non-colocalized genes and nearest genes respectively (Fig. 5 bottom, Supplementary Fig. 7 and 8). We observed a positive correlation between xQTL –log(P-value) and successful target enrichment which is likely due to the increased confidence in the xQTL and therefore a more robust relationship between gene expression and genotype (Supplementary Fig. 9). Lastly, we found that gene and or tissue specificity of PICCOLO genes resulted in a greater enrichment in both OMIM and Pharmaprojects analyses (Fig. 5), suggesting such evidence may be relevant in assessing a gene’s biological relevance.

In summary, we presented a new method that estimates the colocalization between GWAS and xQTL signals without the need for full summary statistics. This innovation greatly expands the number of GWAS and xQTL datasets that can be tested. We observed that xQTLs of colocalized genes tend to be enriched in biologically relevant tissues and enriched for genes linked to rare disease or targets of approved drugs. This work further supports the observations that targets with genetic evidence are more likely to succeed in the clinic^44^ and highlights the importance of mapping the correct genes to genetic associations. Our findings also provide motivation for additional xQTL studies in novel cell types.

Together, these data offer the most comprehensive evidence to date that colocalization testing identifies potentially causal genes. As such, we anticipate that PICCOLO and other colocalization approaches will improve the identification of drug targets from the growing wealth of omic data.

## Methods

### PICCOLO methodology

PICCOLO is available on github (https://github.com/Ksieber/piccolo) as an R package. This code enables users to download PICS^10^ credible sets from the BROAD Institute website (https://pubs.broadinstitute.org/pubs/finemapping/pics.php) and then test two credible sets for colocalization. We removed all credible sets that returned only a single SNP entry from PICS because PICS is unable to differentiate between credible sets with single causal SNPs and SNPs missing from the imputation panel used for the PICs tool. Less than 3% of index SNPs were removed for this reason. The colocalization code is an adaptation from Giambartolomei *et. al*^5^. The default priors are set to be consistent with the default coloc code, where the prior of either genetic signal is 1 × 10^−4^, and the prior of two genetic signals being shared is 1 × 10^−5^.

### xQTLs

Top-hit xQTLs were readily available from sources outlined in Supplementary Table 1. For a given cell type/tissue in each study, we defined the index SNP for each gene as the SNP with the lowest p-value. Using these xQTL index SNPs, we computed the PICS credible sets as outlined in *Farh et al*. (https://pubs.broadinstitute.org/pubs/finemapping/pics.php). Since the cell types and tissues were from multiple sources under different stimuli, we manually grouped cell types into broader categories like those used by the GTEx consortium (Supplementary Table 3).

### GWAS

Index SNPs from the NHGRI-EBI GWAS Catalog were downloaded on November 22^nd^ 2017 from https://www.ebi.ac.uk/gwas/downloads. Using these GWAS index SNPs, we computed the PICS credible sets using the same methods used for the xQTL datasets mentioned above. To create a more consistent naming convention for GWAS traits, each trait and was manually mapped to a MeSH term using the MeSH browser, and subsequently assigned to one of 18 trait categories using a similar methodology outlined in Nelson *et al*^44^. Some traits did not match up with a particular MeSH term or category and were classified as “Miscellaneous” (Supplementary Data 3).

### Calculating PICCOLO systematically

For every GWAS credible set that we generated, we determined all genes within 500 MB of the index GWAS SNP. For each gene within the 500 kb window, we determined every gene-tissue combination of available xQTL credible sets and calculated the PICCOLO score for every gene-tissue xQTL and GWAS combination.

### Calibration with coloc

First, we investigated the distribution of PICCOLO H4 > 0 scores and observed a bimodal distribution (Supplementary Fig. 1). Next, we compared PICCOLO with coloc using a tissue specific approach where the two test results are compared for every trait, in every gene, in every tissue. By evaluating the test statistics of PICCOLO (Supplementary Figure 2), we determined that a PICCOLO cutoff of H4 ≥ 0.9 yields strong predictive power of coloc while maintaining reasonable sensitivity (Supplemental Fig. 2b, positive predictive value = 0.88, true positive rate = 0.45). Given that PICCOLO is estimating many factors, we next asked whether PICCOLO is better at identifying the correct gene at each locus regardless of the tissue specificity. Using this tissue agnostic approach, we observed that PICCOLO H4 ≥ 0.9 was again ideal and yielded a dramatic improvement in the sensitivity compared to the tissue specific approach (Fig. 2b, Supplementary Fig. 3), positive predictive value = 0.89, true positive rate = 0.52). Lastly, given that PICCOLO is not able to access the uncertainty of the strength of association for the two genetic signals (coloc H0, H1, & H2), we hypothesized that PICCOLO would benefit from titrating eQTL P-value cutoffs. Using the tissue agnostic approach, we determined that a 1 × 10^−5^ eQTL P-value threshold is ideal predicting coloc results (Supplementary Fig. 3 c, d).

### Tissue enrichments

A gene was classified as “shared” if at least 1 eQTL for that gene colocalized with GWAS traits within more than one disease category. To measure the enrichment of tissue groups across each trait category, we first identified all colocalizations with genes that were unique to a single trait category and calculated the proportion of those colocalizations contributed by each tissue. We then calculated the proportion of contribution of each tissue for genes that were shared (i.e. not unique to a single category). Statistical differences in tissue proportion in category specific vs shared genes across tissues was tested using Fisher’s exact test (fisher.test) in R.

### Gene enrichment for rare diseases and successful drugs

Rare disease genes were downloaded from the OMIM catalog (https://www.omim.org/) accessed June 5^th^,2018 (Supplementary Data 5). The catalog of successful drug targets was obtained from Pharmaprojects accessed August 5^th^, 2017 (https://pharmaintelligence.informa.com/products-and-services/data-and-analysis/pharmaprojects). Each indication was manually mapped MeSH terms and disease categories. Indications that did not fit into a category were annotated as “Miscellaneous”. A gene was considered a successful target if there was an approved drug in the United State or European Union that targeted it. Due to the proprietary nature of the gene-indication pairs within the Pharmaproject data, we are unable to share the specific list used here in the supplemental information. However, we included a supplemental table specifying the number of successful drugs per MeSH and the number of targets with PICCOLO evidence. Moreover, we repeated our successful target analysis using the publicly available target-indication data from Nelson *et. al*^44^. and observed nearly identical results (Supplementary Fig. 10).

To test for enrichments, we constructed a 2 × 2 contingency table of genes present in GENCODE (v17) or RefSeq (v37.1). Each box was populated with counts based on the presence or absence of the gene as a successful target (or rare disease gene) versus the presence or absence of a positive PICCOLO colocalization for that gene. Tissue-specific PICCOLO genes were defined as genes within loci where the colocalized xQTLs were all from a single tissue group. Gene-specific PICCOLO genes were defined as genes within loci where only a single gene was colocalized. Tissue and gene specific loci were defined as those within loci where a single gene was colocalized and the corresponding xQTL(s) were from a single tissue group. Odds ratios and 95% confidence intervals calculated using fisher.test in R. Statistical methods used to calculate enrichments are the same as those published in Nelson *et. al*^44^.

To determine the percentage of successful trait-indication pairs with evidence of colocalization estimated by PICCOLO, we first selected trait indication-pairs that had an approved drug in the United States or European Union. Next, we removed all target-indication pairs with MeSH terms that did not have at least 1 genome-wide significant association in the GWAS catalog from the analysis (Supplementary Data 6). A target-indication pair was annotated as positive if the target gene matched the colocalized gene and the drug indication was in the same trait category as the GWAS trait. 95% confidence intervals were calculated using the normal approximation method.

## Supplementary Figures

**Supplementary Figure 1:**
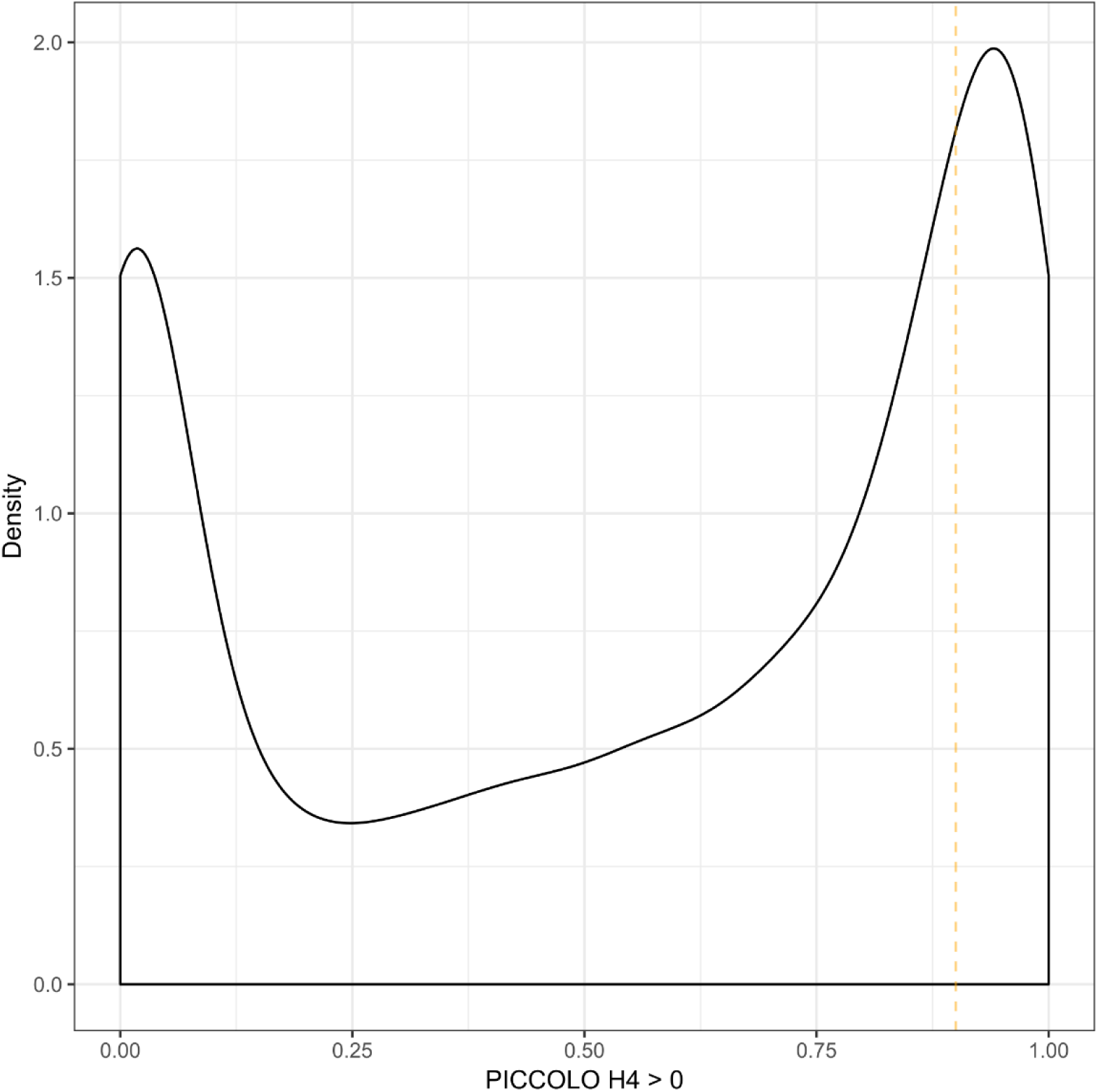
PICCOLO H4 Distribution. Distribution of non-zero PICCOLO H4 scores for associations with 13 traits from the Elliot *et. al*. analysis of the UKBiobank^39^ colocalized with GTEx.

**Supplementary Figure 2:**
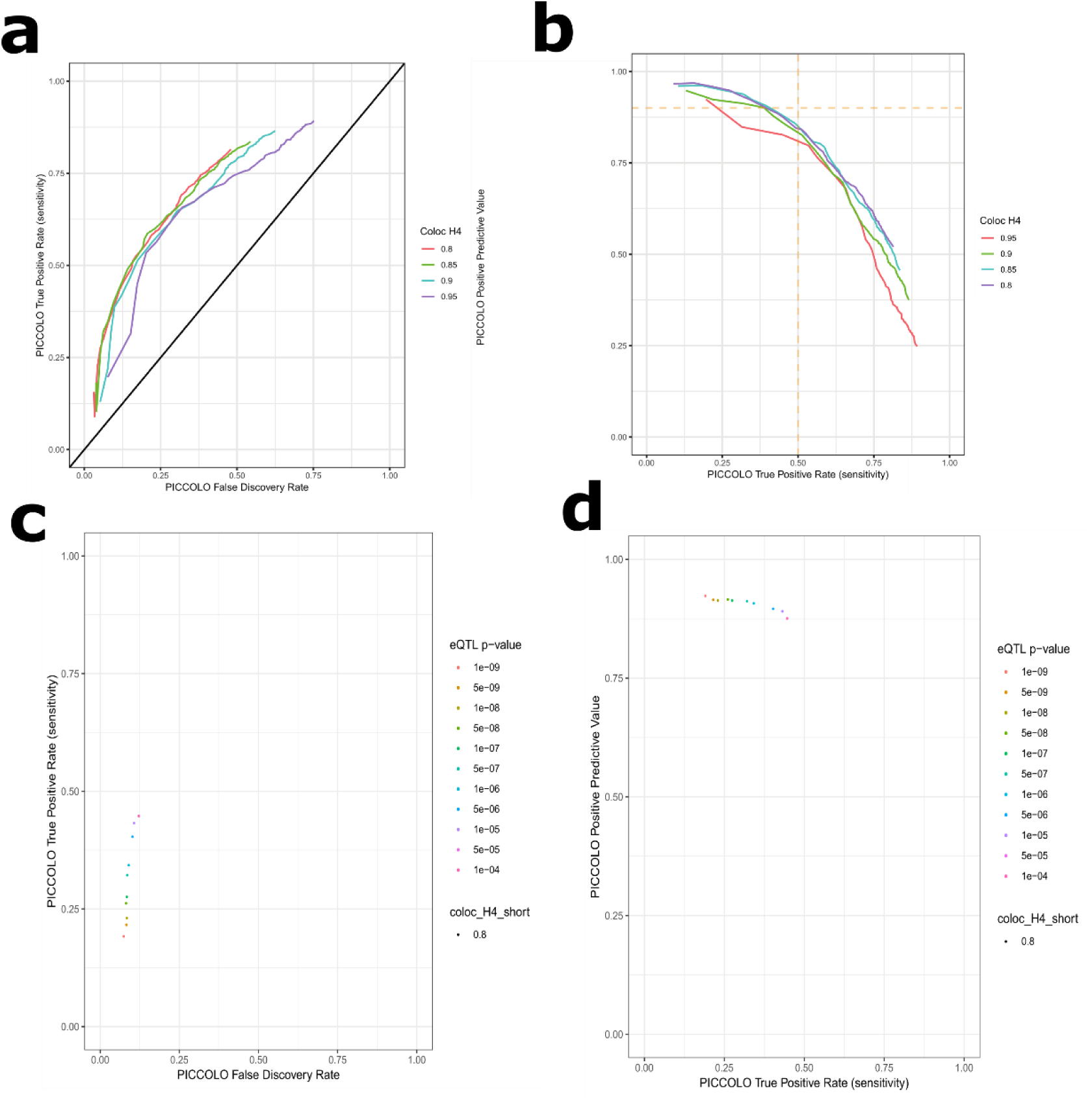
PICCOLO sensitivity tissue specific. **(a)** Using the tissue specific approach, a ROC compares the sensitivity (y-axis) to the false discovery rate (x-axis) of PICCOLO predicting colocalization at multiple coloc H4 thresholds (colored lines). **(b)** The positive predictive value (y-axis) compared the true positive rate (x-axis) of PICCOLO predicting colocalization across multiple coloc H4 thresholds (colored lines). Similar to panels (a) and (b), panels **(c)** and **(d)** compare the same test statistics, respectively, while holding the PICCOLO H4 (0.9) and coloc H4 (0.8) constant while titrating the xQTL P-values (colored dots).

**Supplementary Figure 3:**
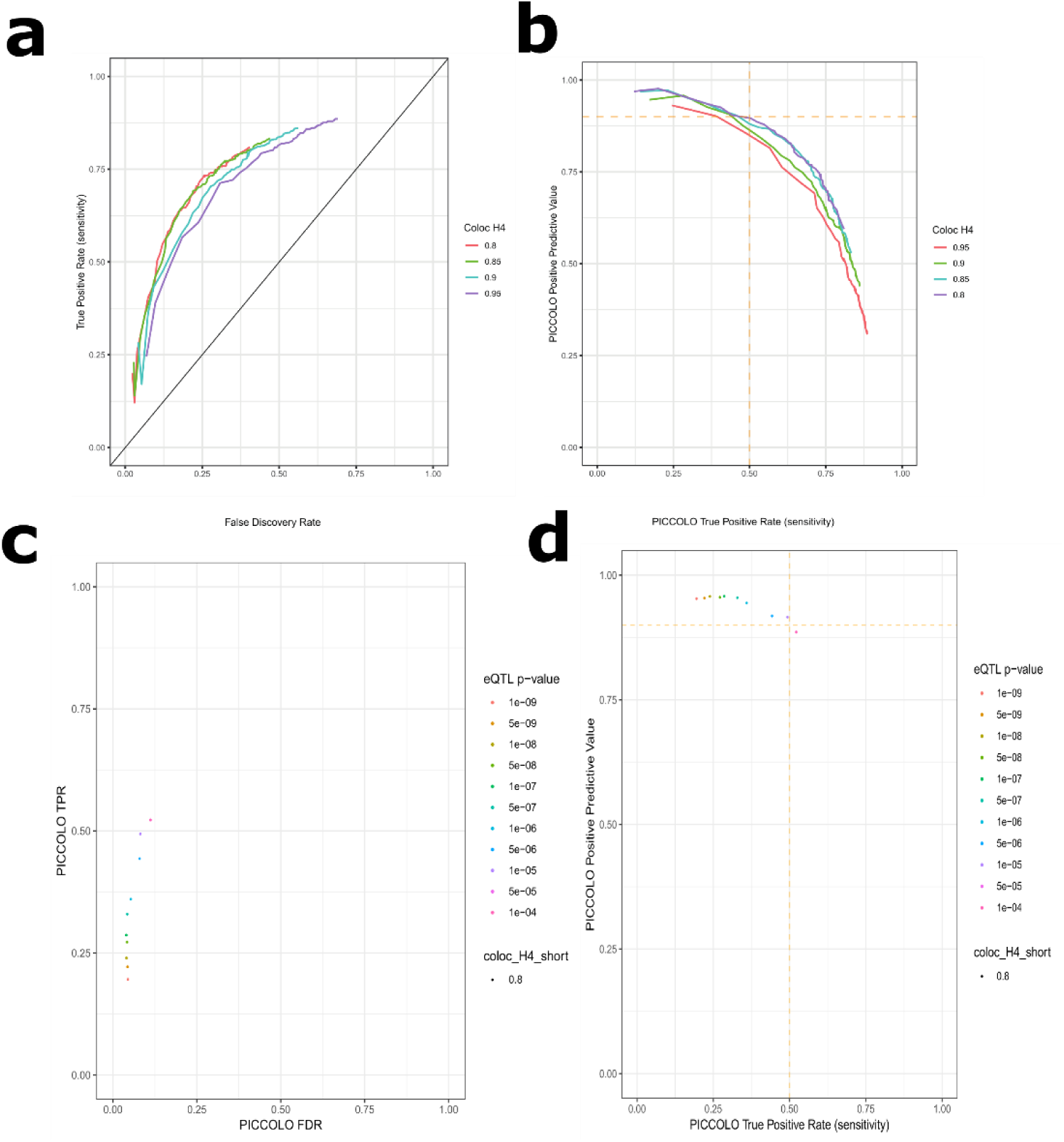
PICCOLO sensitivity tissue agnostic. **(a)** Using the tissue agnostic approach, a ROC compares the sensitivity (y-axis) to the false discovery rate (x-axis) of PICCOLO predicting colocalization at multiple coloc H4 thresholds (colored lines). **(b)** The positive predictive value (y-axis) compared the true positive rate (x-axis) of PICCOLO predicting colocalization across multiple coloc H4 thresholds (colored lines). Similar to panels (a) and (b), panels **(c)** and **(d)** compare the same test statistics, respectively, while holding the PICCOLO H4 (0.9) and coloc H4 (0.8) constant while titrating the xQTL P-values (colored dots).

**Supplementary Figure 4:**
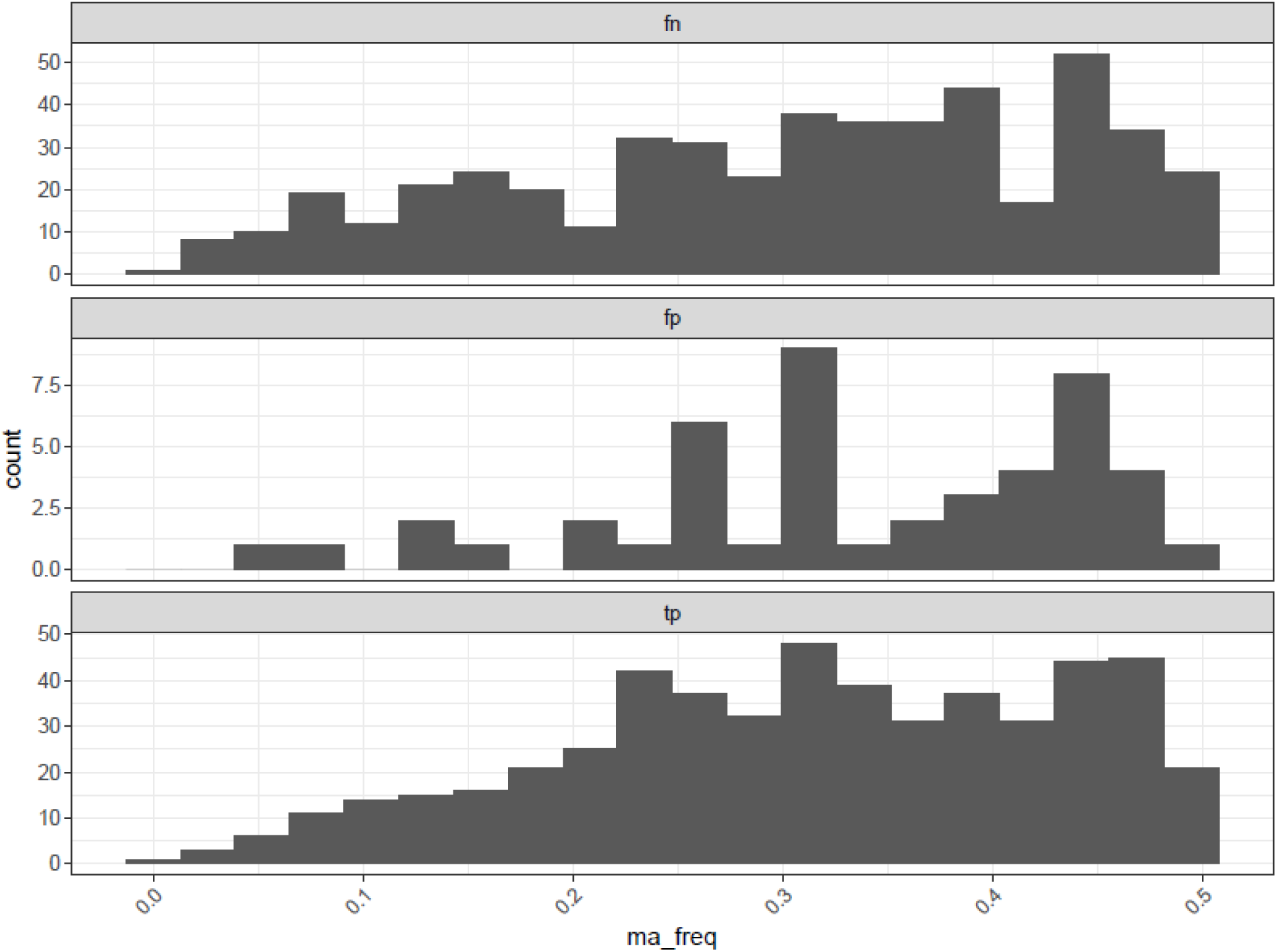
The distribution of the tested GWAS minor allele frequency index SNPs used to compare PICCOLO and coloc. Using the PICCOLO and coloc comparison dataset, PICCOLO results were determined to be either false negatives (fn, top), false positives (fp, mid), or true positives (tp, bottom) based on the results of coloc (H4 ≥ 0.80). For each category, the distribution (y-axis) of the minor allele frequency (x-axis) of the GWAS index SNP is plotted.

**Supplementary Figure 5:**
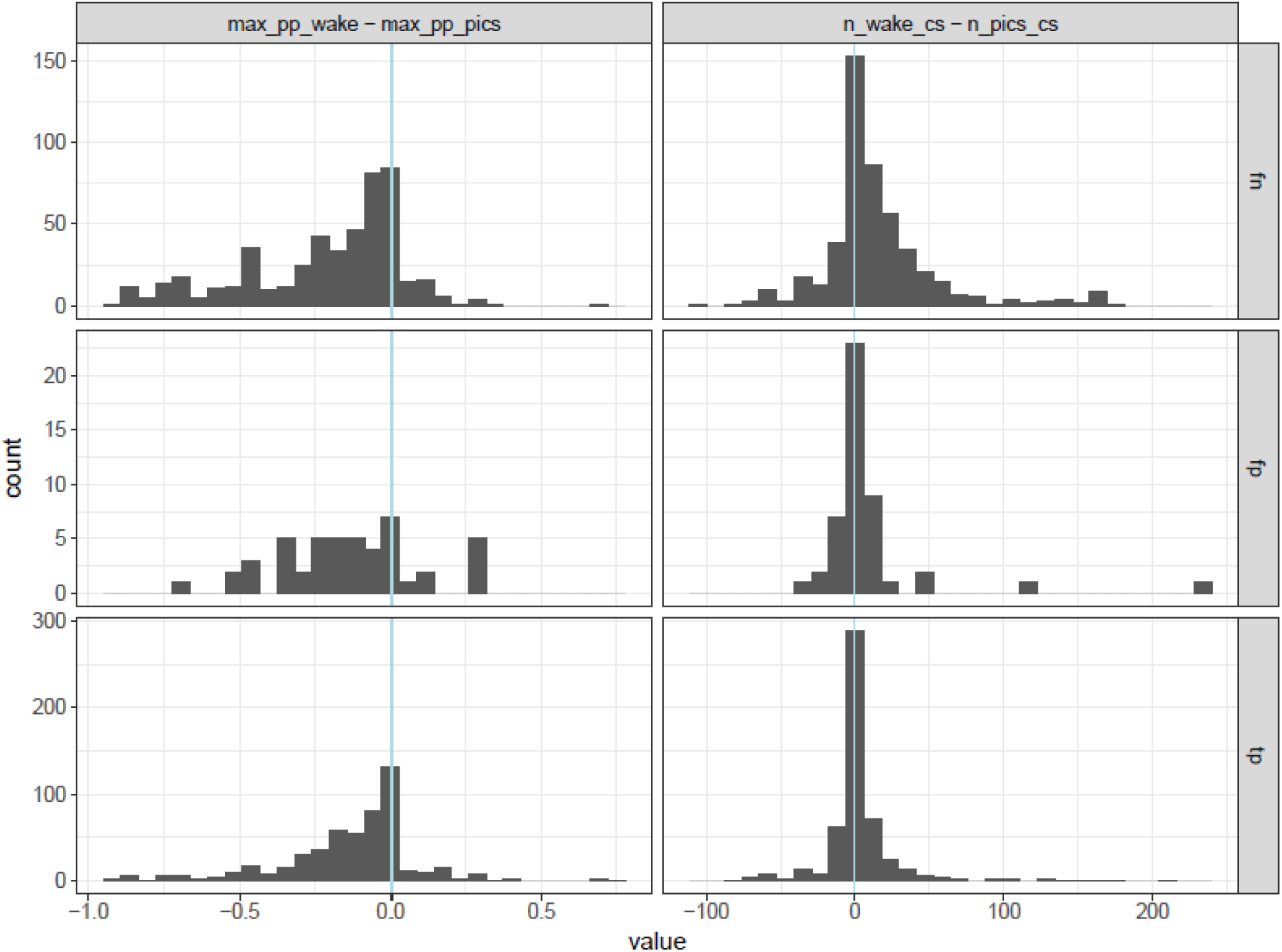
Comparison of the Wakefield and PICS GWAS credible set size and max posterior probability. Using the PICCOLO and coloc comparison dataset, PICCOLO results were determined to be either false negatives (fn, top), false positives (fp, mid), or true positives (tp, bottom) based on the results of coloc (H4 ≥ 0.80). For each category, the distributions of the difference between Wakefield (used for coloc) and the PICs max posterior probability (left panels) and number of SNPs (right panels) for each GWAS credible set are plotted. The vertical blue line illustrates the count where there were no differences between the two methods.

**Supplementary Figure 6:**
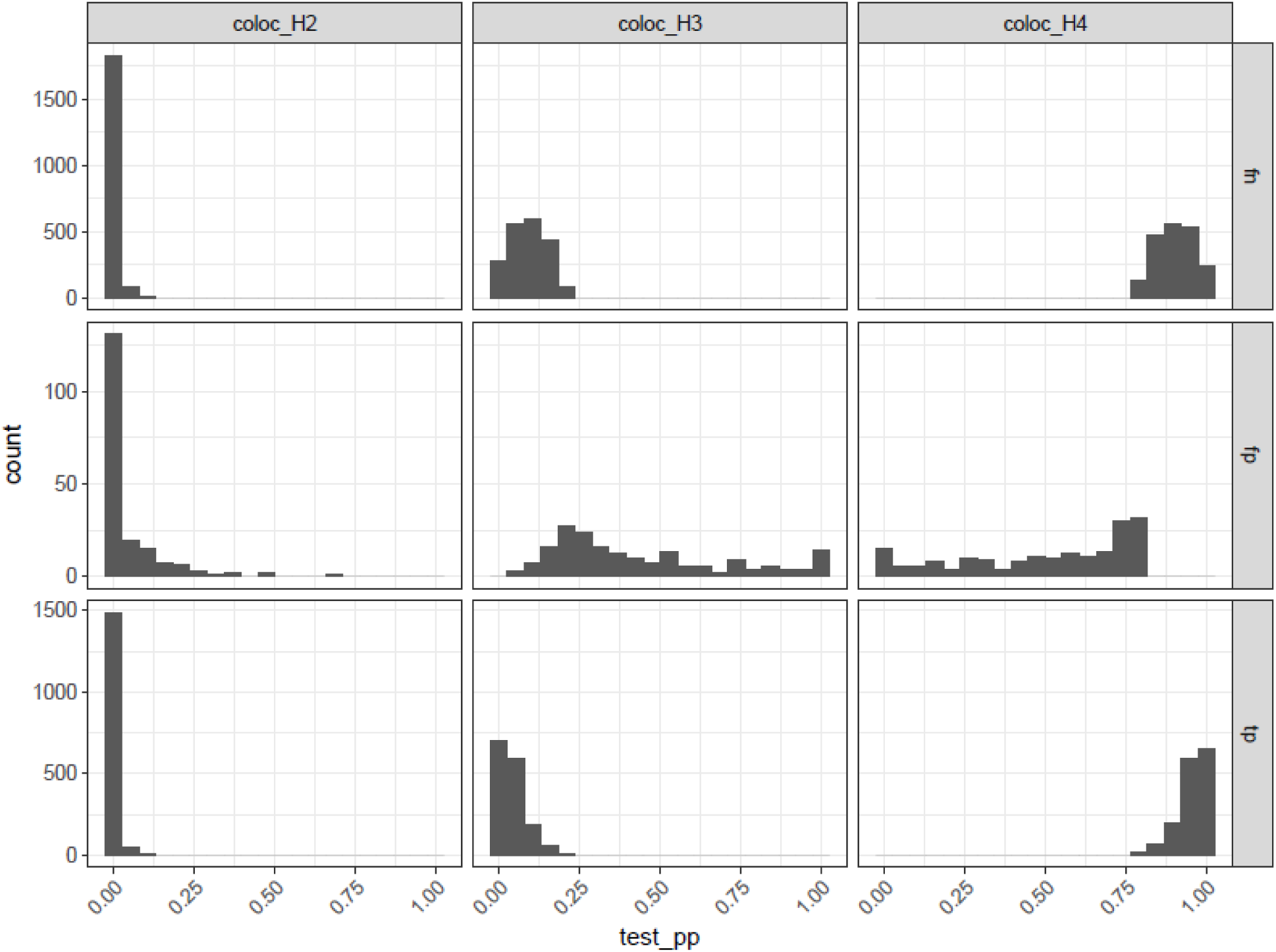
distribution of differences in H2, H3, and H4 for false positives, and true positives. Using the PICCOLO and coloc comparison dataset, PICCOLO results were determined to be either false negatives (fn, top), false positives (fp, mid), or true positives (tp, bottom) based on the results of coloc (H4 ≥ 0.80). For each category, the distribution (y-axis) of the alternative coloc hypotheses test posterior probabilities (x-axis, test_pp) are plotted.

**Supplementary Figure 7:**
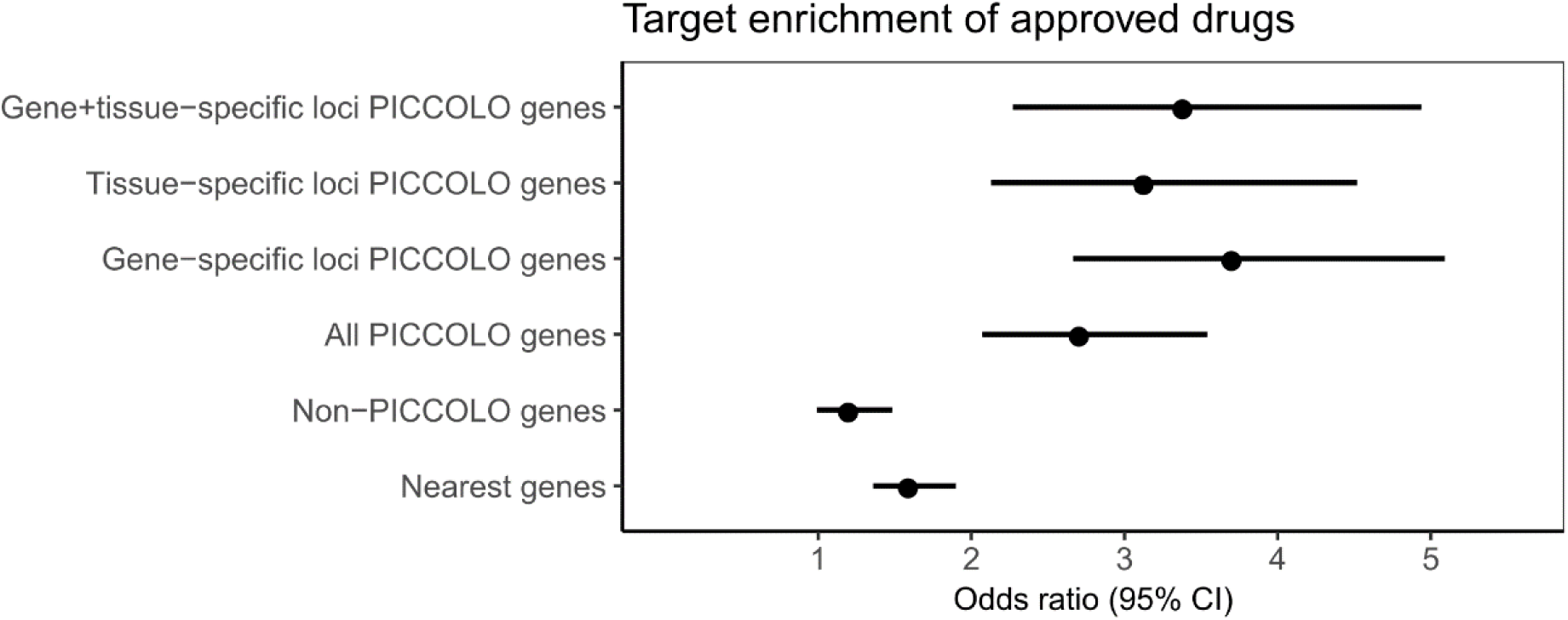
Enrichment of approved drug targets using a PICCOLO H4 ≥ 0.9 and an QTL P ≤ 1 × 10-5. Enrichment was is tested for genes nearest to GWAS index SNPs (Nearest genes), genes that were found to not be colocalized using PICCOLO (Non-PICCOLO genes), all genes found to be colocalized using PICCOLO (All PICCOLO genes), PICCOLO genes within loci where only one gene is colocalized (Gene-specific loci PICCOLO genes), genes within loci where colocalization occur in a single tissue (Tissue-specific loci PICCOLO genes), and PICCOLO genes within loci where a single gene is colocalized in a single tissue (Gene+tissue-specific loci PICCOLO genes). Bars show 95% confidence intervals.

**Supplementary Figure 8:**
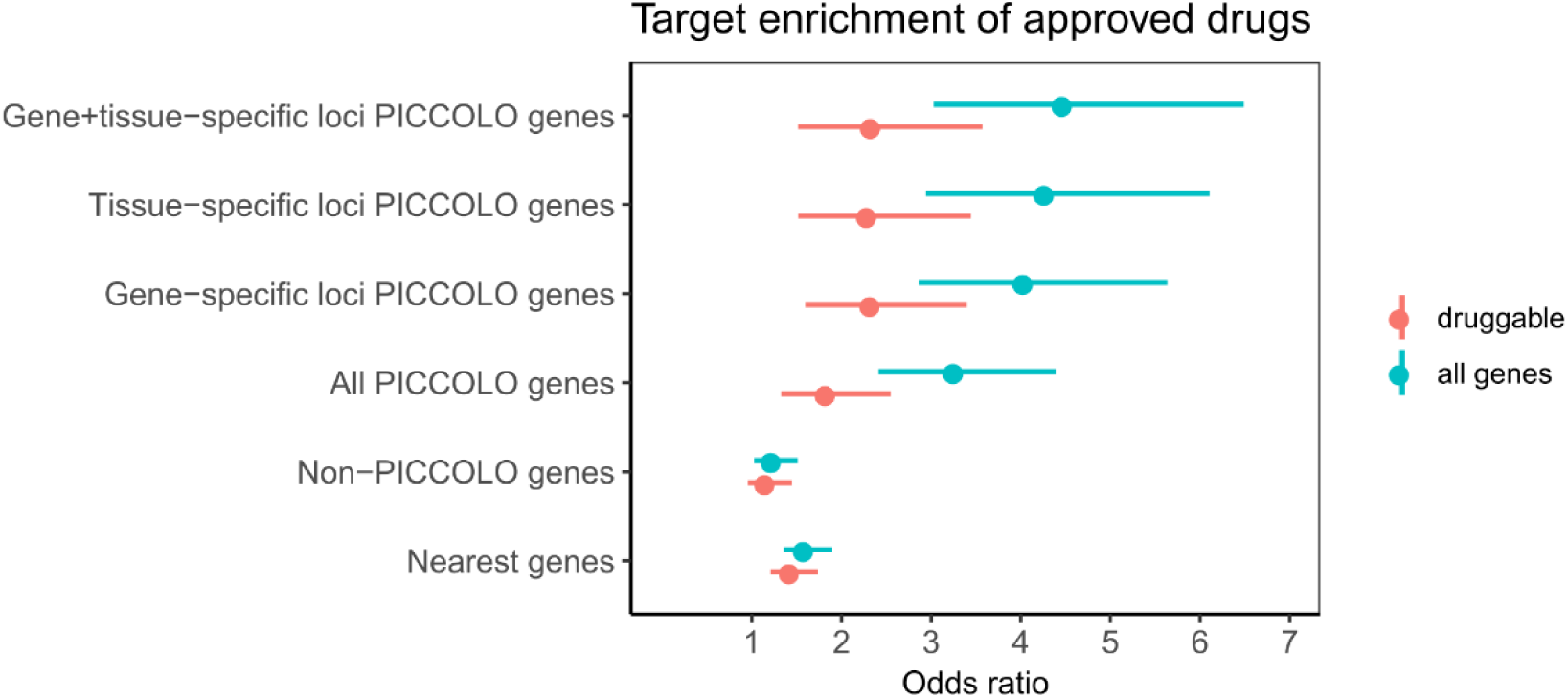
Enrichment of approved drug targets for all genes and the druggable genome. PICCOLO gene enrichment for approved drug targets compared to all protein coding genes in the genome (blue) and all druggable genes (red). Enrichment was is tested for genes nearest to GWAS index SNPs (Nearest genes), genes that were found to not be colocalized using PICCOLO (Non-PICCOLO genes), all genes found to be colocalized using PICCOLO (All PICCOLO genes), PICCOLO genes within loci where only one gene is colocalized (Gene-specific loci PICCOLO genes), genes within loci where colocalization occur in a single tissue (Tissue-specific loci PICCOLO genes), and PICCOLO genes within loci where a single gene is colocalized in a single tissue (Gene+tissue-specific loci PICCOLO genes). Bars show 95% confidence intervals.

**Supplementary Figure 9:**
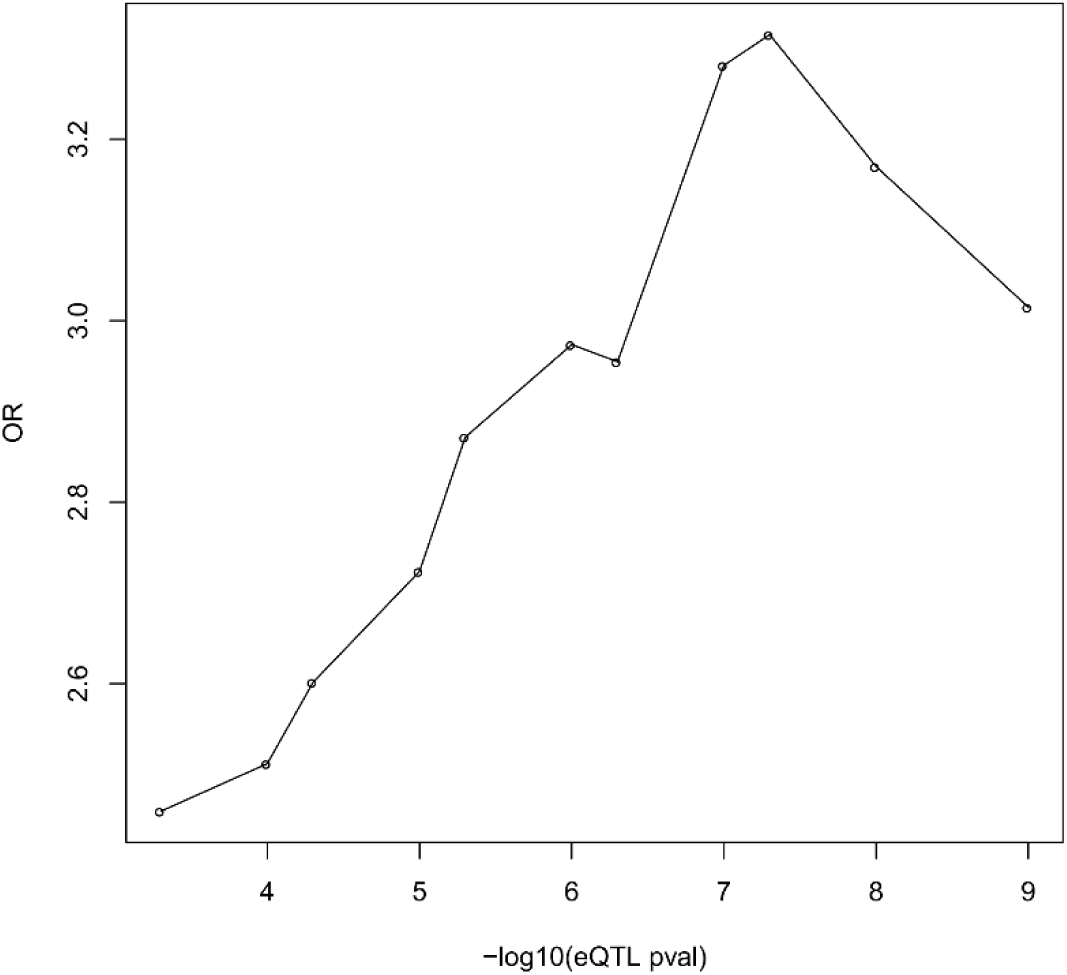
Enrichment of approved drug targets in relation to xQTL P-value cutoff. Points show the mean PICCOLO gene enrichment at xQTL –log10(P-value cutoff) (x-axis).

**Supplementary Figure 10:**
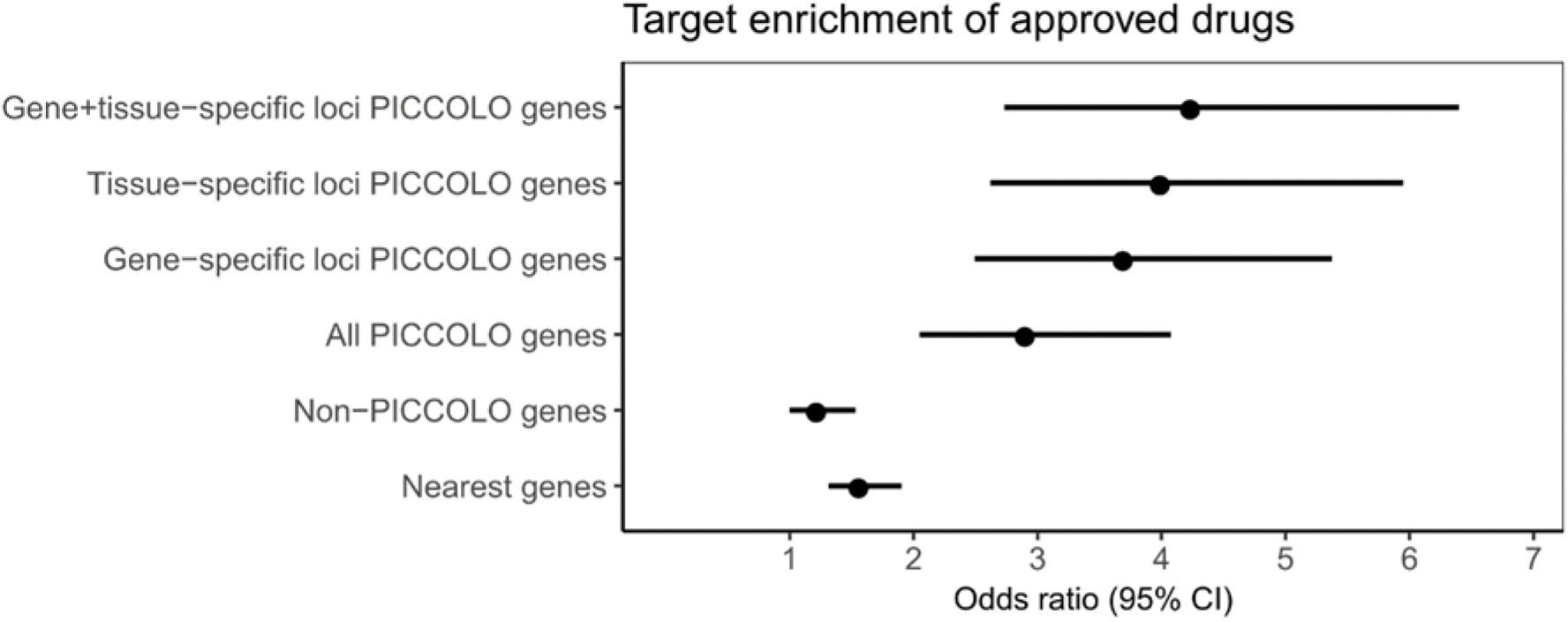
Enrichment of approved drug targets using the Nelson et. al. 2015 successful target-indication pairs dataset. Enrichment was is tested for genes nearest to GWAS index SNPs (Nearest genes), genes that were found to not be colocalized using PICCOLO (Non-PICCOLO genes), all genes found to be colocalized using PICCOLO (All PICCOLO genes), PICCOLO genes within loci where only one gene is colocalized (Gene-specific loci PICCOLO genes), genes within loci where colocalization occur in a single tissue (Tissue-specific loci PICCOLO genes), and PICCOLO genes within loci where a single gene is colocalized in a single tissue (Gene+tissue-specific loci PICCOLO genes). Bars show 95% confidence intervals.

## Supplementary Tables

**Supplementary Table 1:** QTLs sources used in PICCOLO analyses.

| First Author | PMID | TISSUE | QTL type |
| --- | --- | --- | --- |
| GTEX | 29022597 | Multi | eQTL |
| Lee | 24604203 | Dendritic cells (derived from monocytes) | eQTL |
| Barreiro | 22233810 | Dendritic cells | eQTL |
| Fairfax | 24604202 | Monocytes | eQTL |
| Kim-Hellmuth | 28814792 | Monocytes | eQTL |
| Ye | 25214635 | CD4+ T cells | eQTL |
| Nédélec | 27768889 | Monocyte derived Macrophage | eQTL |
| Çalışkan | 25874939 | PBMCs | eQTL |
| Davenport | 26917434 | Leucocytes | eQTL |
| Quach | 27768888 | Monocytes | eQTL |
| Franco | 23878721 | Whole blood | eQTL |
| Naranbhai | 26151758 | Neutrophils | eQTL |
| Raj | 24786080 | CD4(+) T cells and monocytes | eQTL |
| Ferraro | 24610777 | CD4(+) T cells and T regs | eQTL |
| Fairfax | 22446964 | monocytes and B-cells | eQTL |
| Kasela | 28248954 | CD4+ and CD8+ T cells | eQTL |
| Westra | 24013639 | whole blood | eQTL |
| Zhernakova | 27918533 | whole blood | eQTL |
| Hao | 23209423 | lung | eQTL |
| Qui | 21949713 | sputum from COPD patients (ECLIPSE) | pQTL |
| Parisien | 28564610 | Dorsal Root Ganglion | eQTL |
| UBIOPRED | NA | Serum from asthma patients | pQTL |
| Schadt | 18462017 | Liver | eQTL |
| Innocenti | 21637794 | Liver | eQTL |
| Suhre | 28240269 | plasma | pQTL |
| Sun | 29875488 | plasma | pQTL |
| Varshney | 28193859 | islet | eQTL |
| Ko | 28575649 | kidney | eQTL |

**Supplementary Table 2:** GWAS used for comparison between coloc and PICCOLO.

| Analysis | Description | ncase | ncohort |
| --- | --- | --- | --- |
| neale17_1239 | Current tobacco smoking (UKB Broad) | NA | 337030 |
| neale17_1558 | Alcohol intake frequency. (UKB Broad) | NA | 336965 |
| neale17_20002_1111 | Non-cancer illness code, self-reported: asthma (UKB Broad) | 39049 | NA |
| neale17_20002_1112 | Non-cancer illness code, self-reported: chronic obstructive airways disease/copd (UKB Broad) | 1179 | NA |
| neale17_20002_1473 | Non-cancer illness code, self-reported: high cholesterol (UKB Broad) | 41296 | NA |
| neale17_2090 | Seen doctor (GP) for nerves, anxiety, tension or depression (UKB Broad) | 115328 | NA |
| neale17_21001 | Body mass index (BMI) (UKB Broad) | NA | 336107 |
| neale17_47 | Hand grip strength (right) (UKB Broad) | NA | 335842 |
| neale17_50 | Standing height (UKB Broad) | NA | 336474 |
| neale17_6150_4 | Vascular/heart problems diagnosed by doctor: High blood pressure (UKB Broad) | 91033 | NA |
| neale17_6159_4 | Pain type(s) experienced in last month: Back pain (UKB Broad) | 85221 | NA |
| neale17_78 | Heel bone mineral density (BMD) T-score, automated (UKB Broad) | NA | 194398 |
| neale17_I25 | Diagnoses - main ICD10: I25 Chronic ischemic heart disease (UKB Broad) | 8755 | NA |

**Supplementary Table 3:** Tissue grouping key for xQTL studies.

| Tissue group | Tissue type | Cell type |
| --- | --- | --- |
| blood | b-cell | B-cell |
| blood | t-cell | CD4 |
| blood | t-cell | CD4-cis-allconditions |
| blood | t-cell | CD4-meta-cis |
| blood | dendritic | Dendritic-cells-Flu |
| blood | dendritic | Dendritic-cells-IFNb |
| blood | dendritic | Dendritic-cells-LPS |
| blood | dendritic | Dendritic-cells-naive |
| blood | dendritic | Dendritic-cells-re-Flu |
| blood | dendritic | Dendritic-cells-re-IFNb |
| blood | dendritic | Dendritic-cells-re-LPS |
| blood | dendritic | Dendritic-MTB |
| blood | dendritic | Dendritic-naive |
| blood | dendritic | Dendritic-re-MTB |
| lung | Epithelium-airway-cis | Epithelium-airway-cis |
| lung | Epithelium-airway-trans | Epithelium-airway-trans |
| adipose_tissue | gtex-v6p-adipose-subcutaneous | gtex-v6p-adipose-subcutaneous |
| adipose_tissue | gtex-v6p-adipose-visceral-omentum | gtex-v6p-adipose-visceral-omentum |
| adrenal_gland | gtex-v6p-adrenal-gland | gtex-v6p-adrenal-gland |
| blood_vessel | gtex-v6p-artery-aorta | gtex-v6p-artery-aorta |
| blood_vessel | gtex-v6p-artery-coronary | gtex-v6p-artery-coronary |
| blood_vessel | gtex-v6p-artery-tibial | gtex-v6p-artery-tibial |
| bladder | gtex-v6p-bladder | gtex-v6p-bladder |
| brain | gtex-v6p-brain-amygdala | gtex-v6p-brain-amygdala |
| brain | gtex-v6p-brain-anterior-cingulate-cortex-ba24 | gtex-v6p-brain-anterior-cingulate-cortex-ba24 |
| brain | gtex-v6p-brain-caudate-basal-ganglia | gtex-v6p-brain-caudate-basal-ganglia |
| brain | gtex-v6p-brain-cerebellar-hemisphere | gtex-v6p-brain-cerebellar-hemisphere |
| brain | gtex-v6p-brain-cerebellum | gtex-v6p-brain-cerebellum |
| brain | gtex-v6p-brain-cortex | gtex-v6p-brain-cortex |
| brain | gtex-v6p-brain-frontal-cortex-ba9 | gtex-v6p-brain-frontal-cortex-ba9 |
| brain | gtex-v6p-brain-hippocampus | gtex-v6p-brain-hippocampus |
| brain | gtex-v6p-brain-hypothalamus | gtex-v6p-brain-hypothalamus |
| brain | gtex-v6p-brain-nucleus-accumbens-basal-ganglia | gtex-v6p-brain-nucleus-accumbens-basal-ganglia |
| brain | gtex-v6p-brain-putamen-basal-ganglia | gtex-v6p-brain-putamen-basal-ganglia |
| brain | gtex-v6p-brain-spinal-cord-cervical-c-1 | gtex-v6p-brain-spinal-cord-cervical-c-1 |
| brain | gtex-v6p-brain-substantia-nigra | gtex-v6p-brain-substantia-nigra |
| breast | gtex-v6p-breast-mammary-tissue | gtex-v6p-breast-mammary-tissue |
| blood | b-cell | gtex-v6p-cells-ebv-transformed-lymphocytes |
| skin | gtex-v6p-cells-transformed-fibroblasts | gtex-v6p-cells-transformed-fibroblasts |
| cervix_uteri | gtex-v6p-cervix-ectocervix | gtex-v6p-cervix-ectocervix |
| cervix_uteri | gtex-v6p-cervix-endocervix | gtex-v6p-cervix-endocervix |
| colon | gtex-v6p-colon-sigmoid | gtex-v6p-colon-sigmoid |
| colon | gtex-v6p-colon-transverse | gtex-v6p-colon-transverse |
| esophagus | gtex-v6p-esophagus-gastroesophageal-junction | gtex-v6p-esophagus-gastroesophageal-junction |
| esophagus | gtex-v6p-esophagus-mucosa | gtex-v6p-esophagus-mucosa |
| esophagus | gtex-v6p-esophagus-muscularis | gtex-v6p-esophagus-muscularis |
| fallopian_tube | gtex-v6p-fallopian-tube | gtex-v6p-fallopian-tube |
| heart | gtex-v6p-heart-atrial-appendage | gtex-v6p-heart-atrial-appendage |
| heart | gtex-v6p-heart-left-ventricle | gtex-v6p-heart-left-ventricle |
| kidney | gtex-v6p-kidney-cortex | gtex-v6p-kidney-cortex |
| liver | gtex-v6p-liver | gtex-v6p-liver |
| lung | Lung | gtex-v6p-lung |
| salivary_gland | gtex-v6p-minor-salivary-gland | gtex-v6p-minor-salivary-gland |
| muscle | gtex-v6p-muscle-skeletal | gtex-v6p-muscle-skeletal |
| nerve | gtex-v6p-nerve-tibial | gtex-v6p-nerve-tibial |
| ovary | gtex-v6p-ovary | gtex-v6p-ovary |
| pancreas | gtex-v6p-pancreas | gtex-v6p-pancreas |
| pituitary | gtex-v6p-pituitary | gtex-v6p-pituitary |
| prostate | gtex-v6p-prostate | gtex-v6p-prostate |
| skin | gtex-v6p-skin-not-sun-exposed-suprapubic | gtex-v6p-skin-not-sun-exposed-suprapubic |
| skin | gtex-v6p-skin-sun-exposed-lower-leg | gtex-v6p-skin-sun-exposed-lower-leg |
| small_intestine | gtex-v6p-small-intestine-terminal-ileum | gtex-v6p-small-intestine-terminal-ileum |
| spleen | gtex-v6p-spleen | gtex-v6p-spleen |
| stomach | gtex-v6p-stomach | gtex-v6p-stomach |
| testis | gtex-v6p-testis | gtex-v6p-testis |
| thyroid | gtex-v6p-thyroid | gtex-v6p-thyroid |
| uterus | gtex-v6p-uterus | gtex-v6p-uterus |
| vagina | gtex-v6p-vagina | gtex-v6p-vagina |
| blood | blood | gtex-v6p-whole-blood |
| liver | hepatocytes | HLC |
| ipsc | ipsc | IPSC |
| pancreas | islet | Islet |
| kidney | kidney | Kidney |
| blood | leukocyte | Leuco-sepsis |
| liver | liver | Liver |
| lung | lung | Lung-cis |
| blood | macrophage | Macro-Listeria |
| blood | macrophage | Macro-naive |
| blood | macrophage | Macro-re-Listeria |
| blood | macrophage | Macro-re-Salmonella |
| blood | macrophage | Macro-Salmonella |
| blood | monocyte | Mono |
| blood | monocyte | Monocyte |
| blood | monocyte | Mono-Flu |
| blood | monocyte | Mono-IFN |
| blood | monocyte | Mono-LPS |
| blood | monocyte | Mono-LPS-24h |
| blood | monocyte | Mono-LPS-2h |
| blood | monocyte | Mono-LPS-6h |
| blood | monocyte | Mono-LPS-90m |
| blood | monocyte | Mono-mdp-6h |
| blood | monocyte | Mono-mdp-90m |
| blood | monocyte | Mono-naive |
| blood | monocyte | Mono-Pam3CSK4 |
| blood | monocyte | Mono-R848 |
| blood | monocyte | Mono-re-Flu |
| blood | monocyte | Mono-re-LPS |
| blood | monocyte | Mono-re-LPS-6h |
| blood | monocyte | Mono-re-LPS-90m |
| blood | monocyte | Mono-re-mdp-6h |
| blood | monocyte | Mono-re-mdp-90m |
| blood | monocyte | Mono-re-Pam3CSK4 |
| blood | monocyte | Mono-re-R848 |
| blood | monocyte | Mono-re-rna-6h |
| blood | monocyte | Mono-re-rna-90m |
| blood | monocyte | Mono-rna-6h |
| blood | monocyte | Mono-rna-90min |
| blood | neutrophil | Neutro |
| blood | blood | PBMC-naive |
| blood | blood | PBMC-re-rv |
| blood | blood | PBMC-rv |
| plasma | plasma | Plasma-pQTL |
| plasma | plasma | Serum-cis-pQTL |
| plasma | plasma | Serum-pQTL-cis |
| plasma | plasma | Serum-trans-pQTL |
| lung | sputum | Sputum-COPD-patients-ECLIPSE-cis |
| blood | t-cell | Treg |
| blood | blood | WB-naive |

## Supplementary Data Sets

Supplementary Data Set 1: PICCOLO and coloc results for 13 traits colocalized with GTEx

Supplementary Data Set 2: All PICCOLO results

Supplementary Data Set 3: GWAS Catalog with MeSH annotations

Supplementary Data Set 4: Number of tissues and diseases per gene

Supplementary Data Set 5: OMIM Genes

Supplementary Data Set 6: Number of associations per MeSH

Supplementary Data Set 7: Number of successful drug targets with PICCOLO evidence by MeSH

## Author contributions

C.G. and K.B.S jointly conceptualized the study and lead the analyses. J.E.G. and M.R.H obtained the xQTL datasets and performed the MeSH annotations. T.J. and M.R.N provided scientific guidance and assisted with the analyses. C.G. and K.B.S wrote the manuscript with assistance from all other authors.

## Acknowledgements

We would like to thank Meg G. Ehm for her suggestions on the manuscript. We are appreciative of John H. Riley, Stewart A. Bates, and the rest of the U-BIOPRED group for allowing us to use their unpublished pQTL results.

## Competing interests

All authors are employees of GlaxoSmithKline.

